# Gene Program Negotiation Defines Cellular Identity in Single-Cell Transcriptomes

**DOI:** 10.64898/2026.07.05.736629

**Authors:** Ji-Yong Sung, Jae-Ho Cheong

**Affiliations:** Department of Neurosurgery, Seoul National University Bundang Hospital, Seoul National University College of Medicine, Seongnam-si, Republic of Korea; Department of Surgery, Yonsei University College of Medicine, Seoul 03722, Republic of Korea; Department of Quantum Information Science, Graduate School, Yonsei University, Seoul 03722, Republic of Korea

## Abstract

Single-cell transcriptomics has transformed the characterization of cellular heterogeneity by enabling systematic analysis of biological gene programs. However, existing computational approaches primarily quantify the activity of individual programs independently and therefore provide limited insight into how multiple simultaneously active programs collectively determine cellular identity. Here we present Gene Program Negotiation (GPN), a graph-based computational framework that models regulatory decision-making among concurrently active biological programs. GPN reconstructs cell-specific program interaction networks from local transcriptional neighborhoods and quantifies regulatory organization using the Gene Program Coherence Index (GPCI) together with measures of local regulatory conflict, program diversity, and dominance. These graph-derived properties enable the classification of individual cells into five regulatory decision states: Consensus, Competition, Negotiation, Dominance, and Low activity. Applying GPN to gastric cancer single-cell transcriptomes revealed that cells sharing the same dominant biological program frequently occupied distinct regulatory decision states, demonstrating that dominant program identity alone does not uniquely define cellular regulatory organization. Competition states consistently exhibited elevated local regulatory conflict and were preferentially enriched among transition-like cells, indicating that regulatory competition is closely associated with transcriptional plasticity. Independent validation using glioblastoma single-cell transcriptomes reproduced these regulatory patterns without modification of the computational framework, supporting the robustness and generalizability of the approach across biologically distinct malignancies. These findings establish regulatory negotiation as an additional layer of cellular organization beyond conventional gene-program activity analysis. By explicitly modeling interactions among simultaneously active biological programs, GPN provides a general computational framework for investigating regulatory coordination, cellular plasticity, and dynamic cell-state organization in single-cell transcriptomic data.

## Introduction

The emergence of single-cell RNA sequencing (scRNA-seq) has fundamentally transformed our understanding of cellular heterogeneity by enabling transcriptomic profiling at single-cell resolution.^1,2^ Recent advances in computational analysis have enabled the identification of distinct cell populations, developmental trajectories, regulatory programs, and disease-associated cellular states across diverse biological systems.^3^ Gene program analysis, in particular, has become one of the most widely used approaches for interpreting single-cell transcriptomes by summarizing coordinated expression of biologically related genes into functional modules. Despite these advances, current computational frameworks primarily quantify the activity of individual gene programs independently.^4,5^ Widely used methods, including module-score approaches, gene set enrichment analysis, and latent-factor models, estimate whether a specific biological process such as stemness, epithelial–mesenchymal transition (EMT), proliferation, immune activation, or stress response is active within a cell. ^6,7^ However, these methods provide limited information regarding how multiple regulatory programs collectively determine cellular identity.^8,9^ This limitation is particularly important because biological cells rarely activate only a single regulatory program.^10^ Tumor cells frequently exhibit simultaneous activation of stemness and EMT programs, immune cells integrate inflammatory and metabolic signals,^11–16^ and stromal cells coordinate multiple extracellular matrix and signaling pathways.^12,17–24^ The final transcriptional state of a cell therefore emerges from the integration of multiple concurrent regulatory processes rather than from any individual program alone.^25,26^ Consequently, two cells with similar program activity profiles may nevertheless occupy distinct regulatory states depending on how these programs interact.^27,28^ Current single-cell analysis lacks a computational framework capable of explicitly modeling this regulatory integration process.^9^ Existing approaches generally describe what biological programs are active but do not address how these programs cooperate, compete, or dominate one another to establish a stable cellular phenotype.^5^ As a result, an important layer of regulatory organization underlying cellular decision-making remains largely unexplored. ^28,29^

Here, we propose Gene Program Negotiation (GPN), a computational framework that models regulatory decision-making as an interaction process among simultaneously active gene programs. Rather than treating gene programs as independent measurements, GPN considers each active program as contributing a weighted regulatory influence that interacts with other programs within the same cell.^5,30^ These interactions are integrated into a cell-specific regulatory negotiation network, enabling quantitative characterization of cooperative and antagonistic regulatory relationships among biological processes.^31^ To summarize the overall organization of regulatory interactions, we introduce the Gene Program Coherence Index (GPCI), a quantitative metric describing the degree of coordinated regulatory decision-making within individual cells. In addition, GPN classifies cells into distinct regulatory decision states—including Consensus, Competition, Dominance, Negotiation, and Low activity—according to the balance between regulatory coherence, conflict, program dominance, and overall regulatory activity. We demonstrate the utility of GPN using publicly available gastric cancer single-cell transcriptomic data. Compared with conventional gene program analyses, GPN reveals an additional layer of regulatory organization that cannot be inferred from program activity scores alone.^32^ By explicitly modeling interactions among simultaneously active regulatory programs, our framework provides a quantitative description of cell-specific regulatory decision-making and offers a general computational strategy for studying cellular state transitions across diverse biological systems.

## Results

### A Gene Program Negotiation framework for modeling regulatory decision-making in single cells

Current single-cell transcriptomic analyses primarily quantify the activity of individual gene programs to characterize cellular states. Although these approaches have greatly improved the interpretation of cellular heterogeneity, they generally treat each gene program as an independent entity. Consequently, they provide limited information about how multiple regulatory programs collectively contribute to the establishment of a single cellular phenotype. In biological systems, however, multiple regulatory programs are frequently activated simultaneously. For example, stemness, epithelial–mesenchymal transition (EMT), stress response, proliferation, and immune-associated programs may coexist within the same cell. The final cellular phenotype therefore emerges not from the activity of a single program but from the combined influence of multiple interacting regulatory programs. Existing single-cell methods do not explicitly model this regulatory integration process. To address this limitation, we developed Gene Program Negotiation (GPN), a computational framework that models the regulatory decision-making process occurring among simultaneously active gene programs (**Fig. 1**). Rather than considering gene program activities independently, GPN assumes that each active regulatory program contributes a weighted regulatory vote toward determining the transcriptional state of an individual cell. These votes are integrated through a cell-specific negotiation process that captures both cooperative and antagonistic interactions among active programs. Within this framework, each cell is represented by a cell-specific regulatory negotiation network, in which nodes correspond to active gene programs and edges represent local cooperative or competitive regulatory relationships inferred from neighboring cells in transcriptional space. The resulting negotiation network is summarized by the Gene Program Coherence Index (GPCI), which quantitatively measures the degree of coordinated regulatory decision-making within each cell. Cells with high GPCI exhibit coherent agreement among multiple regulatory programs, whereas low GPCI indicates conflicting regulatory influences and unstable transcriptional organization. Based on regulatory coherence, local program conflict, regulatory dominance, and overall program activity, GPN further classifies individual cells into five regulatory decision states: Consensus, Competition, Dominance, Negotiation, and Low activity (**Fig. 1**). Unlike conventional approaches that describe only the magnitude of gene program activation, GPN provides an explicit computational representation of how multiple regulatory programs collectively determine cellular identity. This framework establishes a quantitative basis for investigating regulatory decision-making across heterogeneous cell populations and can be applied to diverse single-cell transcriptomic datasets to characterize dynamic cellular states beyond conventional gene program activity analysis.

**Figure 1.**
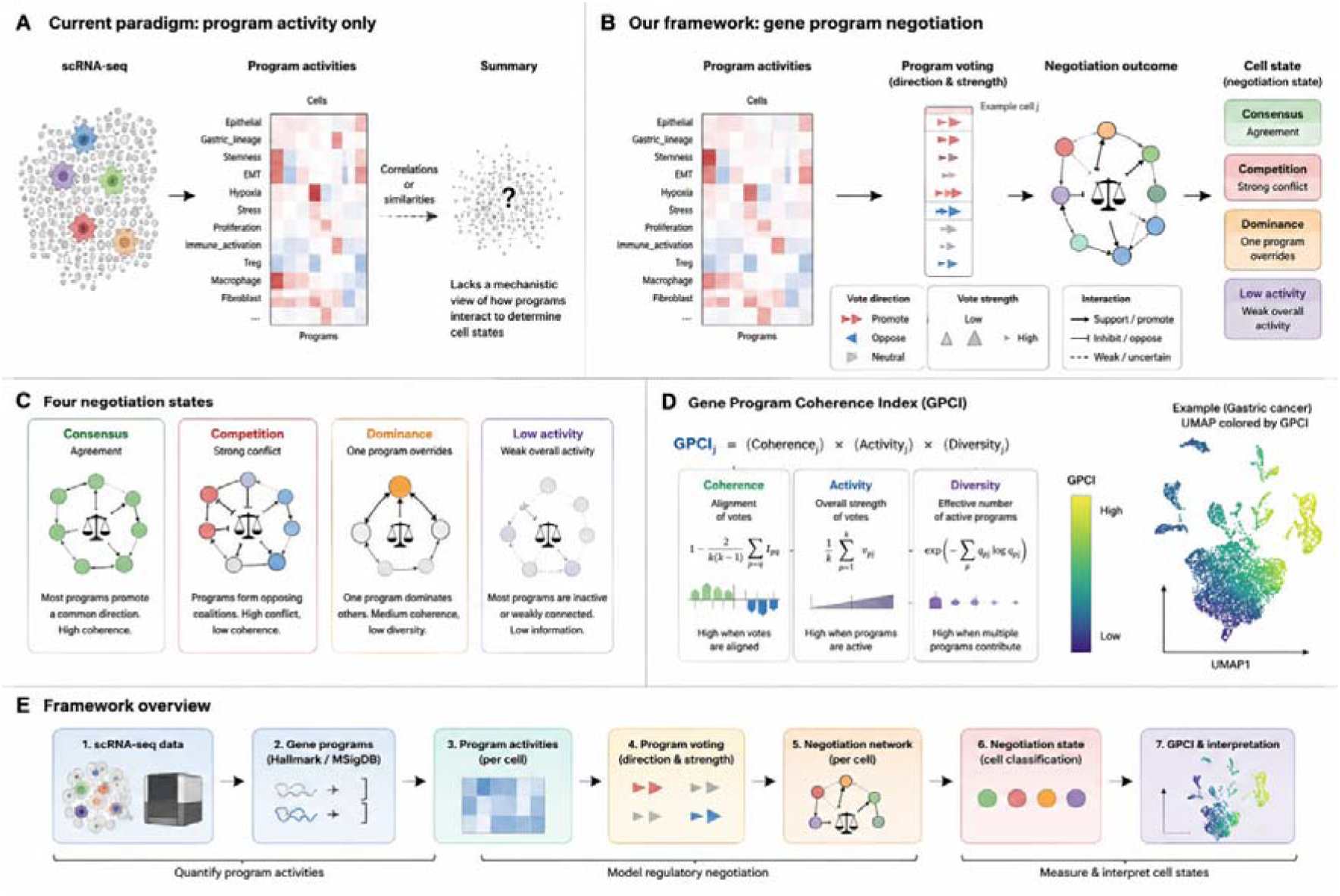
Gene Program Negotiation framework for quantifying regulatory decision-making in single cells. (A) Current single-cell transcriptomic analyses summarize cells by independent gene program activities, providing limited information about how multiple programs collectively determine cellular identity. Although individual program scores can be quantified, interactions among simultaneously active programs are not explicitly modeled. (B) Overview of the proposed Gene Program Negotiation (GPN) framework. Gene program activities are first quantified for each cell using predefined or data-driven regulatory programs. Each active program subsequently casts a directional vote representing its regulatory influence (promoting, suppressing, or neutral) with a corresponding voting strength. The collection of votes is integrated into a cell-specific negotiation network, which models cooperative and competitive interactions among active gene programs. The negotiation outcome determines the regulatory state of each individual cell. (C) Conceptual illustration of the five regulatory decision states identified by the framework. Consensus represents coordinated agreement among multiple gene programs. Competition reflects simultaneous activation of opposing regulatory programs. Negotiation represents balanced integration among multiple simultaneously active biological programs without complete dominance or antagonism. Dominance indicates that a single program overwhelmingly determines the regulatory state. Low activity corresponds to cells with weak or sparse regulatory activity. (D) Definition of the Gene Program Coherence Index (GPCI). GPCI integrates three complementary components: (i) coherence, measuring the consistency of regulatory votes among active programs; (ii) activity, representing the overall regulatory strength of active programs; and (iii) diversity, quantifying the effective number of participating gene programs. Higher GPCI values indicate coordinated regulatory decision-making, whereas lower values indicate conflicting or incoherent program activities. (E) Overall computational workflow of the Gene Program Negotiation framework. Starting from single-cell RNA sequencing data, gene program activities are estimated, transformed into regulatory votes, assembled into a negotiation network for each cell, summarized by GPCI, and finally interpreted to characterize regulatory decision states across the cellular population.

### Gene Program Negotiation reveals cell-specific regulatory decision states in gastric cancer single-cell transcriptomes

To evaluate the biological utility of the proposed Gene Program Negotiation (GPN) framework, we applied the method to a publicly available gastric cancer single-cell RNA sequencing dataset comprising tumor, stromal, and immune cell populations. Following quality control, dimensionality reduction, and unsupervised clustering, cells formed well-defined transcriptional compartments corresponding to epithelial/tumor-like, fibroblast/stromal-like, and immune/myeloid-like populations (**Fig. 2A**). We next quantified the activities of twelve biologically relevant regulatory programs, including epithelial identity, gastric lineage, stemness, epithelial–mesenchymal transition (EMT), hypoxia, stress response, proliferation, interferon signaling, immune activation, regulatory T-cell, macrophage, and fibroblast programs. As expected, these programs exhibited distinct activity patterns across Leiden clusters, indicating substantial regulatory heterogeneity within the gastric cancer microenvironment (**Fig. 2B**).

**Figure 2.**
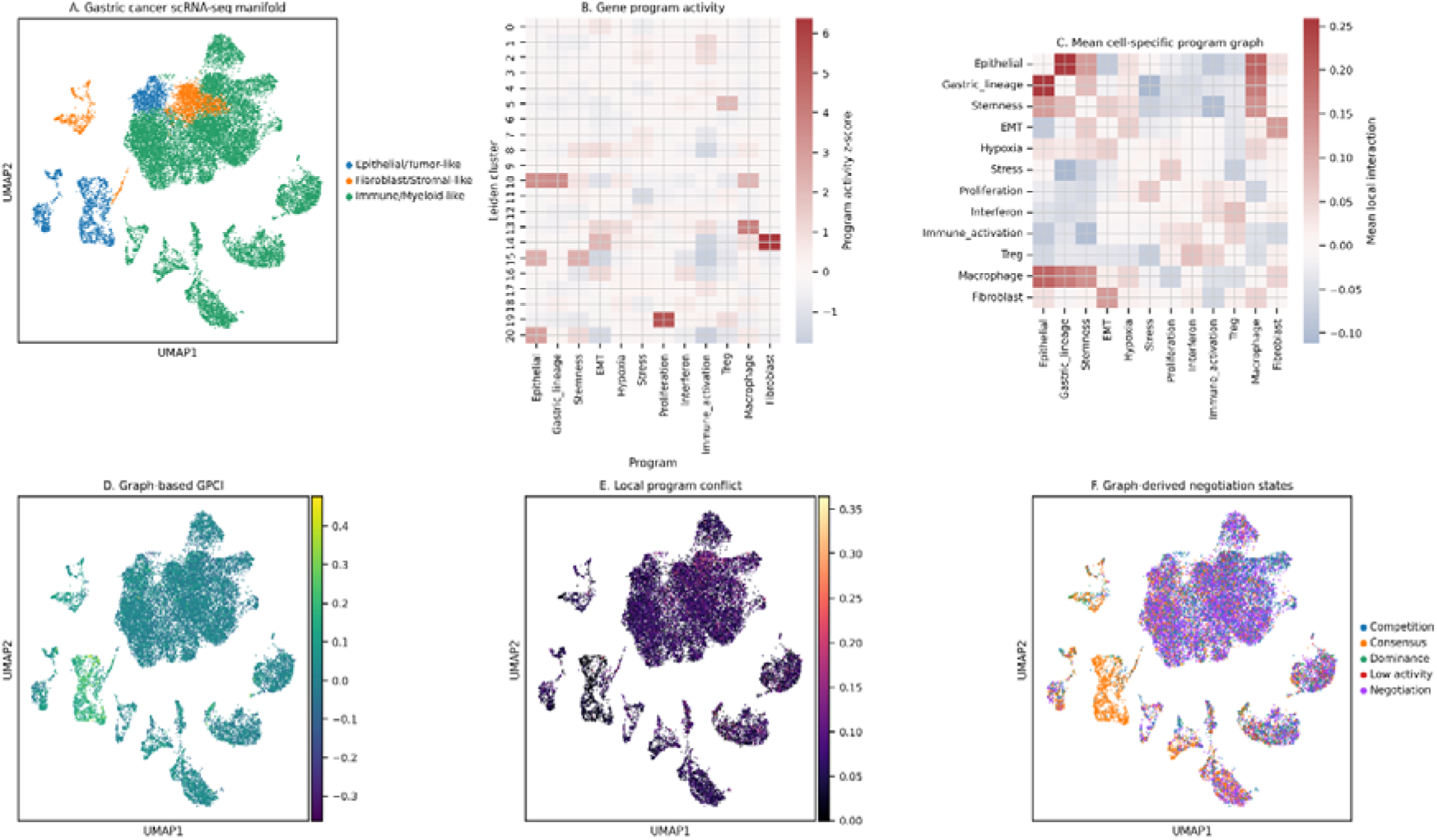
Gene Program Negotiation identifies regulatory decision states in gastric cancer single-cell transcriptomes. (A) UMAP visualization of gastric cancer single-cell transcriptomes showing three major cellular compartments inferred from dominant regulatory programs, including epithelial/tumor-like, fibroblast/stromal-like, and immune/myeloid-like populations. (B) Heatmap of average gene program activities across Leiden clusters. Program activities were quantified using predefined regulatory gene sets representing epithelial identity, gastric lineage, stemness, epithelial–mesenchymal transition (EMT), hypoxia, stress response, proliferation, interferon signaling, immune activation, regulatory T cells, macrophages, and fibroblasts. Values represent standardized program activity scores. (C) Average cell-specific regulatory negotiation network. For each cell, local interactions among active gene programs were estimated from neighborhood-specific regulatory relationships, and the resulting cell-specific negotiation graphs were averaged across all cells. Positive interactions indicate coordinated regulatory programs, whereas negative interactions indicate locally conflicting regulatory activities. (D) UMAP colored by the Gene Program Coherence Index (GPCI). Higher GPCI values indicate coordinated activation of multiple regulatory programs, whereas lower values indicate incoherent or fragmented regulatory decision-making. (E) UMAP showing the local program conflict score for individual cells. High conflict scores identify cells in which simultaneously active regulatory programs exhibit strong local antagonistic interactions, suggesting unstable or transitional regulatory states. (F) Cell-specific negotiation states inferred by the proposed framework. Each cell was classified into one of five regulatory decision states (Consensus, Competition, Dominance, Negotiation, or Low activity) according to the balance between regulatory coherence, conflict, program activity, and dominance. These states summarize distinct modes of regulatory decision-making across the gastric cancer cellular landscape.

Unlike conventional analyses that consider each program independently, GPN estimated local interactions among simultaneously active programs for every individual cell. Averaging these cell-specific negotiation networks revealed characteristic patterns of coordinated and antagonistic regulatory relationships among biological programs (**Fig. 2C**). Epithelial and gastric-lineage programs showed strong cooperative interactions, whereas immune-associated programs displayed regulatory relationships distinct from epithelial programs. These observations suggest that regulatory programs are organized into structured interaction networks rather than functioning independently. To quantify the overall coordination of regulatory programs, we calculated the Gene Program Coherence Index (GPCI) for each cell. GPCI values varied substantially across the cellular landscape, demonstrating that regulatory coherence differs among individual cells despite similar transcriptional identities (**Fig. 2D**). Cells exhibiting higher GPCI values were characterized by stronger agreement among active regulatory programs, whereas lower GPCI values reflected less coordinated regulatory organization. We further quantified local program conflict by measuring antagonistic interactions within each cell-specific negotiation network. Elevated conflict scores were detected in discrete cellular regions, indicating the presence of cells simultaneously influenced by competing regulatory programs (**Fig. 2E**). These cells are consistent with transcriptionally heterogeneous or transitional regulatory states that cannot be identified using program activity scores alone. Finally, integrating regulatory coherence, conflict, dominance, and overall program activity enabled the classification of each cell into one of five regulatory decision states: Consensus, Competition, Dominance, Negotiation, and Low activity (**Fig. 2F**). These regulatory decision states were broadly distributed across all major cellular compartments rather than being restricted to a single transcriptional cluster, suggesting that regulatory decision-making represents an additional layer of biological organization beyond conventional cell-type annotation. Collectively, these results demonstrate that GPN captures cell-specific regulatory organization that is not represented by individual gene program activities alone. By explicitly modeling cooperative and competitive interactions among simultaneously active regulatory programs, the proposed framework provides a quantitative description of regulatory decision-making at single-cell resolution.

### Gene Program Negotiation reveals regulatory organization beyond conventional gene program activity analysis

We next investigated whether the regulatory decision states identified by Gene Program Negotiation (GPN) could be explained using conventional gene program activity analysis. Current single-cell approaches generally assign each cell to its highest-scoring biological program, implicitly assuming that the dominant program adequately represents the regulatory state of the cell. In contrast, GPN explicitly models interactions among simultaneously active gene programs before inferring cell-specific regulatory decision states. To compare these analytical frameworks, we first assigned each cell to its dominant gene program using conventional independent program scoring (**Fig. 3A**). This representation identified the predominant biological process within individual cells but did not capture the regulatory relationships among co-activated programs. Applying GPN to the same dataset revealed a distinct organization of the transcriptomic landscape, classifying cells into five regulatory decision states (Competition, Consensus, Dominance, Negotiation, and Low activity) according to regulatory coherence and local program interactions (**Fig. 3B**). Importantly, each dominant gene program contained multiple negotiation states rather than a single uniform regulatory state (**Fig. 3C**). For example, cells sharing identical dominant programs were distributed across Consensus, Competition, Negotiation, Dominance, and Low activity states with different proportions. These findings indicate that dominant program identity alone does not uniquely determine the underlying regulatory organization of a cell. Consistent with this observation, the Gene Program Coherence Index (GPCI) exhibited substantial variability within individual dominant program categories (**Fig. 3D**). Cells assigned to the same dominant biological program frequently displayed markedly different coherence values, demonstrating that similar program identities can arise from distinct patterns of regulatory coordination. Thus, GPCI captures an additional dimension of regulatory organization that is not represented by conventional dominant-program assignment. We next examined local regulatory conflict across GPN-derived decision states. As expected, Competition cells displayed the highest levels of local program conflict, whereas Consensus cells exhibited consistently low conflict (**Fig. 3E**). Negotiation and Dominance states occupied intermediate conflict levels, suggesting that these regulatory states represent distinct modes of integrating concurrent biological programs rather than arbitrary subdivisions of transcriptional activity. Finally, we evaluated whether regulatory coherence could be explained simply by the magnitude of gene program activation. Maximum program activity showed only a weak association with GPCI (Spearman’s r = 0.11; **Fig. 3F**), indicating that highly active programs do not necessarily correspond to highly coordinated regulatory states. Cells with comparable levels of program activity frequently exhibited markedly different coherence values and belonged to different negotiation states. These observations demonstrate that regulatory decision-making depends primarily on the organization of interactions among concurrently active gene programs rather than on the activity of any single program.

**Figure 3.**
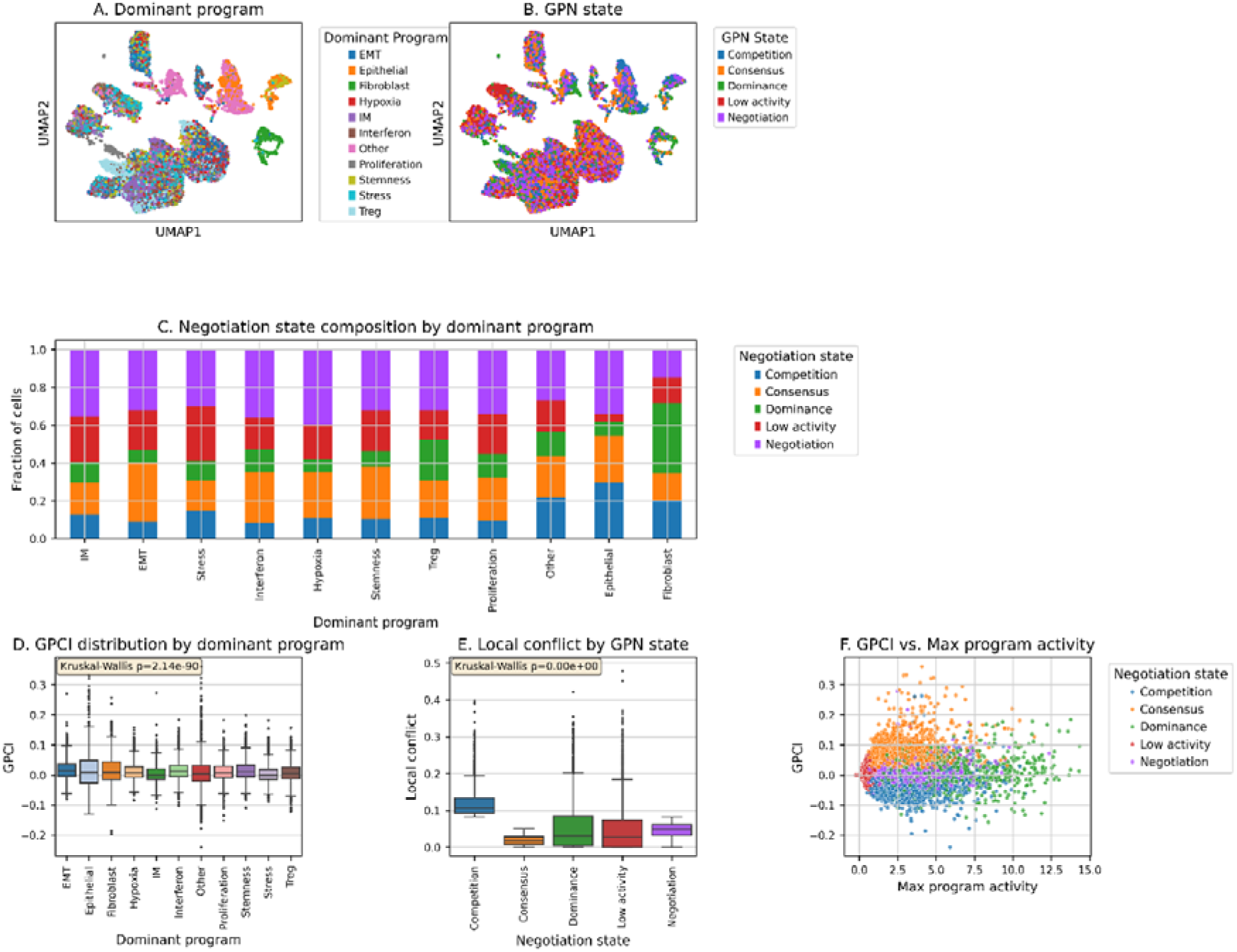
Gene Program Negotiation reveals regulatory information beyond conventional gene program activity analysis. (A) UMAP visualization of gastric cancer single-cell transcriptomes colored according to the dominant gene program assigned to each cell using conventional independent gene program scoring. Cells are labeled by the highest-scoring regulatory program without considering interactions among simultaneously active programs. (B) UMAP visualization of the same dataset colored by Gene Program Negotiation (GPN)-derived regulatory decision states. Unlike conventional program assignment, GPN classifies cells into Competition, Consensus, Dominance, Negotiation, and Low activity states by integrating regulatory coherence, local conflict, program dominance, and overall program activity. (C) Distribution of GPN-derived regulatory decision states within each dominant gene program. Multiple negotiation states are observed within individual dominant program categories, demonstrating that cells sharing the same dominant biological program can occupy distinct regulatory decision states. (D) Distribution of the Gene Program Coherence Index (GPCI) across dominant gene program categories. Considerable heterogeneity in GPCI is observed within individual dominant programs, indicating that regulatory coherence cannot be explained solely by dominant program identity. (E) Local program conflict across GPN-derived regulatory decision states. Cells classified as Competition exhibit substantially higher local conflict than other regulatory decision states, consistent with the presence of competing regulatory programs within individual cells. (F) Relationship between maximum gene program activity and GPCI. The weak correlation between program activity and regulatory coherence demonstrates that the magnitude of gene program activation alone is insufficient to explain regulatory organization at the single-cell level.

Unlike Consensus and Competition, which represent the two extremes of regulatory agreement and antagonism, respectively, Negotiation cells occupied an intermediate regulatory regime characterized by sustained activation of multiple biological programs without overwhelming dominance by any single program. Rather than representing a residual category, these cells exhibited coordinated contributions from multiple regulatory programs while maintaining intermediate levels of graph coherence and local conflict. This organization is consistent with a regulatory balancing process in which multiple biological influences coexist without complete consensus or strong antagonism. Therefore, the Negotiation state represents a distinct mode of regulatory organization that cannot be explained solely by dominant program identity.

Collectively, these results demonstrate that conventional gene program analysis identifies which biological program is most active, whereas GPN additionally quantifies how multiple active programs are coordinated within individual cells. By explicitly modeling regulatory interactions, GPN uncovers an additional layer of cell-state organization that cannot be inferred from gene program activity alone.

### Regulatory decision states are dynamically remodeled during cell-state transition

Having established that GPN identifies regulatory organization beyond conventional gene program activity analysis, we next investigated whether these regulatory decision states are associated with dynamic cellular state transitions. We therefore reconstructed the continuous transcriptional trajectory of gastric cancer cells using diffusion pseudotime and examined how GPN-derived regulatory features evolved along the inferred trajectory. Diffusion pseudotime revealed a continuous organization of the gastric cancer single-cell transcriptomic landscape, capturing progressive transitions among cellular states (**Fig. 4A**). Mapping GPN-derived regulatory decision states onto the same trajectory demonstrated that Competition, Consensus, Dominance, Negotiation, and Low activity states were broadly distributed throughout the cellular manifold but exhibited distinct spatial organizations along the inferred transition axis (**Fig. 4B**). These observations suggest that regulatory decision states are dynamically remodeled during cell-state progression rather than representing static transcriptional subclasses. We next examined how regulatory coherence changed during trajectory progression. Although the overall trend in the Gene Program Coherence Index (GPCI) varied only modestly across pseudotime, substantial cell-to-cell heterogeneity was observed at every stage of the trajectory (**Fig. 4C**). Cells occupying similar pseudotime positions frequently displayed markedly different coherence values, indicating that progression along the transcriptional trajectory does not uniquely determine the underlying regulatory organization. These findings suggest that regulatory coordination represents an additional dimension of cellular state that is only partially captured by conventional trajectory inference. Analysis of local program conflict further supported this interpretation. Cells exhibiting elevated regulatory conflict were detected throughout the trajectory but were particularly enriched within the Competition state, whereas Consensus cells consistently maintained low levels of local conflict (**Fig. 4D**). These results indicate that transcriptionally similar cells can nevertheless differ substantially in the extent of regulatory competition occurring among simultaneously active gene programs. Consistent with these observations, the composition of GPN-derived regulatory decision states changed progressively across pseudotime (**Fig. 4E**). Rather than remaining constant during trajectory progression, the relative abundance of Competition, Consensus, Dominance, Negotiation, and Low activity states continuously shifted along the inferred developmental axis. This dynamic redistribution suggests that cell-state transitions involve coordinated reorganization of regulatory interactions in addition to gradual transcriptional changes. Finally, we asked whether cells undergoing regulatory transition preferentially occupied specific GPN states. Transition-like cells, defined by elevated local conflict together with multi-program activity and intermediate regulatory coherence, were significantly enriched for Competition and Negotiation states compared with the remaining cell population (Fisher’s exact test, odds ratio = 5.09; **Fig. 4F**). This enrichment indicates that cells experiencing regulatory competition are substantially more likely to occupy transitional cellular states than cells with stable regulatory organization. These findings demonstrate that GPN-derived regulatory decision states are dynamically associated with cellular state transitions in gastric cancer. Whereas conventional trajectory analysis describes continuous changes in transcriptional similarity, GPN provides complementary information by quantifying how interactions among concurrently active gene programs are reorganized during the transition process.

**Figure 4.**
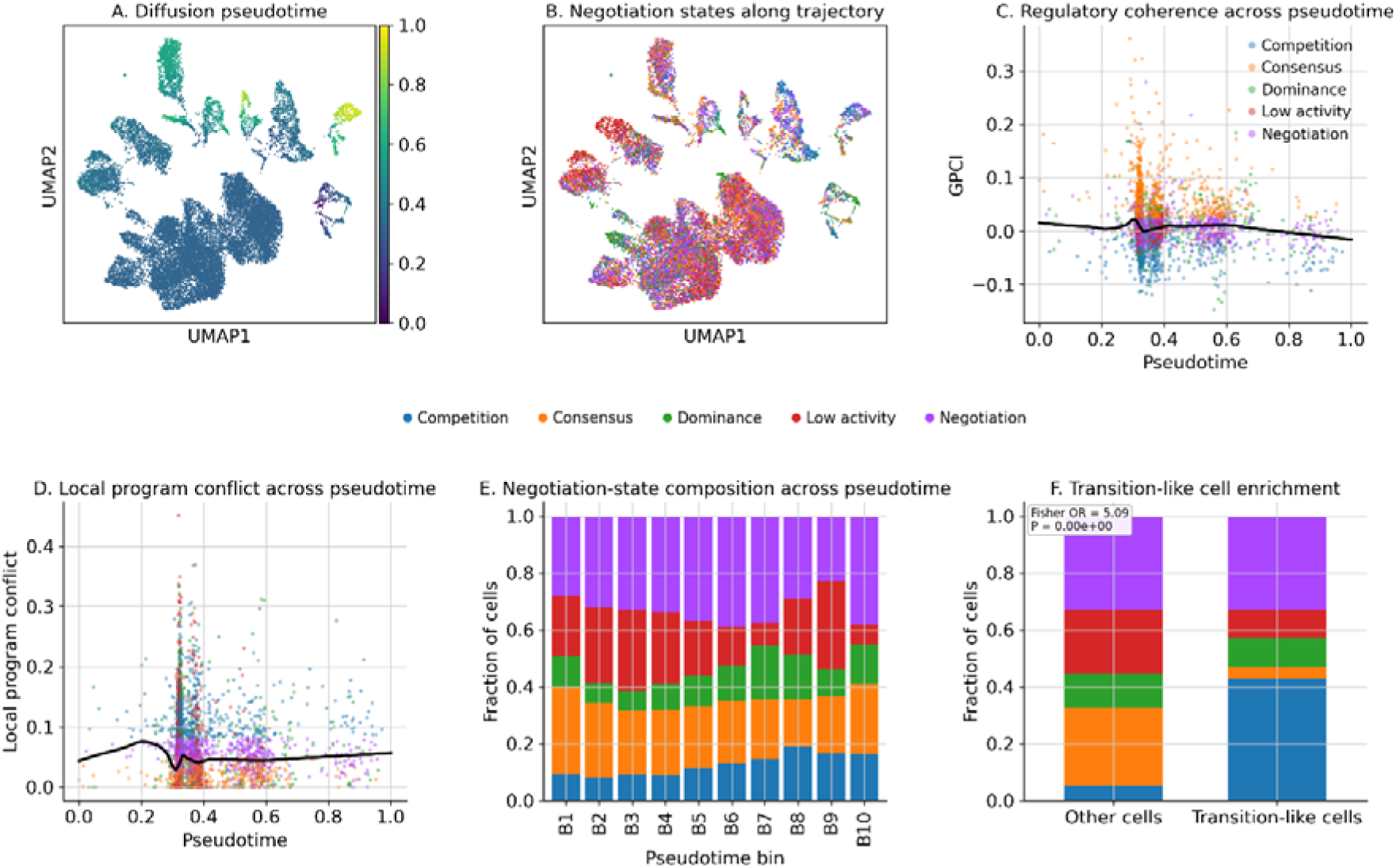
Regulatory decision states are associated with cell-state transition dynamics in gastric cancer single-cell transcriptomes. (A) UMAP visualization of gastric cancer single-cell transcriptomes colored by diffusion pseudotime. Pseudotime was inferred from the transcriptomic manifold using diffusion-based trajectory analysis to reconstruct continuous cell-state progression. (B) Distribution of Gene Program Negotiation (GPN)-derived regulatory decision states along the inferred trajectory. Cells were classified into Competition, Consensus, Dominance, Negotiation, and Low activity states based on regulatory coherence, local program conflict, program dominance, and overall program activity. (C) Relationship between the Gene Program Coherence Index (GPCI) and diffusion pseudotime. Each point represents an individual cell, and the black curve indicates a locally weighted regression (LOESS) fit. GPCI varies continuously along the inferred trajectory, suggesting dynamic changes in regulatory coordination during cell-state progression. (D) Local program conflict as a function of diffusion pseudotime. Competition states exhibit elevated local conflict throughout the trajectory, whereas Consensus cells maintain consistently low levels of regulatory conflict. (E) Relative abundance of GPN-derived regulatory decision states across pseudotime bins. The composition of regulatory decision states changes continuously during cell-state progression, indicating dynamic remodeling of regulatory organization rather than static transcriptional identities. (F) Enrichment of regulatory decision states in transition-like cells. Transition-like cells were operationally defined using combined measures of local program conflict, multi-program activity, and intermediate regulatory coherence. Competition and Negotiation states were significantly enriched within transition-like cells compared with the remaining cell population (Fisher’s exact test, odds ratio = 5.09).

### Biological interpretation of GPN-derived regulatory decision states

To determine whether GPN-derived regulatory decision states represent biologically meaningful cellular organizations rather than purely mathematical classifications, we systematically compared transcriptional programs across all regulatory states. Distinct GPN states exhibited characteristic combinations of gene-program activities, indicating that each state corresponds to a unique regulatory organization (**Fig. 5A**). Consensus cells were characterized by coordinated activation of epithelial, stemness, EMT, hypoxia, stress, and proliferation programs with relatively balanced regulatory interactions. In contrast, Competition cells showed strong activation of stemness- and EMT-associated programs together with elevated hypoxia and stress responses, suggesting simultaneous activation of biologically antagonistic transcriptional modules. Negotiation cells displayed intermediate activity across multiple programs without a single dominant regulatory axis, whereas Dominance cells preferentially activated a limited subset of programs while suppressing alternative transcriptional programs. Low activity cells exhibited uniformly reduced program activity across nearly all regulatory modules. The distribution of GPN states across broad cellular compartments further demonstrated that regulatory decision states were not restricted to a specific lineage (**Fig. 5B**). Instead, epithelial/tumor-like, stromal, and immune-derived cells all contained multiple GPN states, suggesting that regulatory negotiation reflects a general property of cellular transcriptional organization rather than differences in cell identity alone. To investigate whether regulatory decision states were associated with cellular plasticity, we compared transition-like regulatory scores across GPN states (**Fig. 5C**). Competition states displayed the highest transition scores, whereas Consensus and Low activity states showed substantially lower values, indicating that elevated local regulatory conflict is closely associated with increased transcriptional plasticity. Negotiation cells occupied an intermediate position between these extremes, consistent with a transient regulatory decision-making process. We next examined the activity of canonical cancer-associated gene programs within each regulatory state (**Fig. 5D**). Competition and Negotiation states simultaneously exhibited increased activities of Stemness, EMT, Hypoxia, Stress, and Proliferation programs, whereas Dominance states preferentially activated only a restricted subset of these programs. These findings indicate that GPN captures coordinated interactions among multiple regulatory programs rather than variation in individual pathways. To quantify program-specific regulatory preferences, we calculated the enrichment of high program activity within each GPN state using log□odds ratios (**Fig. 5E**). Consensus states were enriched for coordinated activation of multiple compatible programs, whereas Competition states preferentially exhibited enrichment of antagonistic regulatory modules associated with cellular plasticity. In contrast, Low activity states showed uniformly reduced enrichment across nearly all transcriptional programs, supporting their interpretation as globally inactive regulatory states. These analyses demonstrate that GPN-derived regulatory decision states represent biologically distinct modes of transcriptional organization rather than alternative descriptions of dominant gene-program activity (**Fig. 5F**). By integrating local regulatory coherence with simultaneous gene-program activity, GPN identifies conserved regulatory strategies that link mathematical measures of program interaction to interpretable cellular phenotypes.

**Figure 5.**
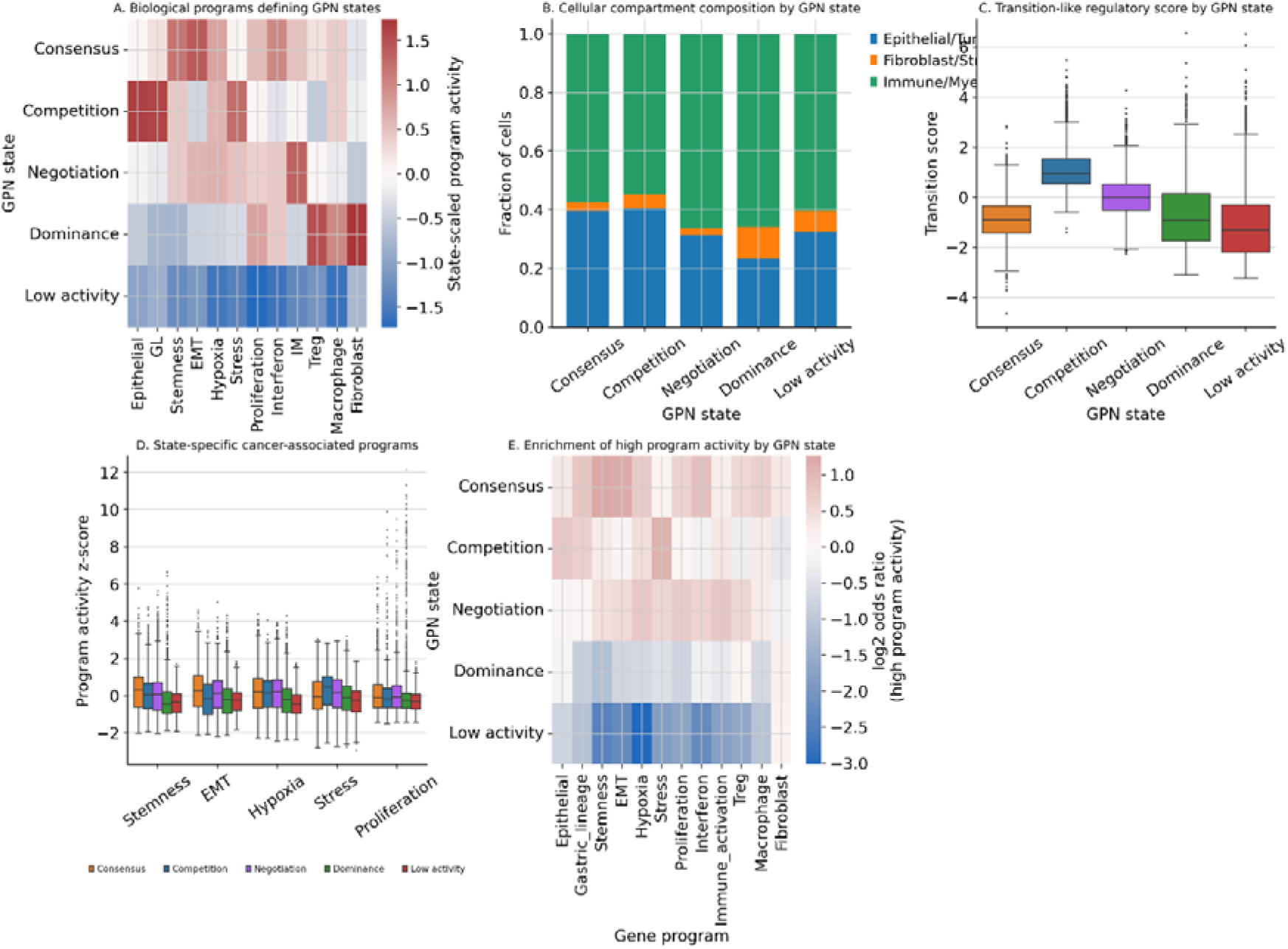
Biological interpretation of Gene Program Negotiation (GPN) states in gastric cancer single cells. (A) Heatmap showing the average activity of representative gene programs across GPN-derived regulatory decision states. Program activities were standardized (z-score) across cells before averaging within each state. Distinct regulatory states exhibit characteristic combinations of coordinated and antagonistic gene programs, indicating that each GPN state corresponds to a unique transcriptional organization rather than differences in a single dominant program. (B) Cellular compartment composition of each GPN state. The relative proportions of epithelial/tumor-like, fibroblast/stromal-like, and immune/myeloid-like cells were calculated for each regulatory decision state, demonstrating that GPN states are observed across multiple cellular compartments and are not restricted to a single lineage. (C) Distribution of transition-like regulatory scores across GPN states. Competition states exhibited the highest transition-like scores, whereas Consensus and Low activity states showed significantly lower values, suggesting that regulatory competition is associated with increased transcriptional plasticity. (D) Distribution of cancer-associated gene program activities (Stemness, EMT, Hypoxia, Stress, and Proliferation) across GPN states. Competition and Negotiation states displayed elevated activities of multiple cancer-associated programs simultaneously, whereas Dominance states preferentially exhibited activation of a restricted subset of programs. (E) Enrichment of high gene-program activity within each GPN state, quantified as the log□odds ratio for cells exhibiting high program activity. Consensus states were enriched for coordinated activation of multiple programs, whereas Competition states preferentially showed enrichment of antagonistic programs associated with cellular plasticity. Low activity states demonstrated uniformly reduced enrichment across all regulatory programs.

### Independent cross-cancer validation demonstrates the generalizability of Gene Program Negotiation

To evaluate whether the regulatory principles identified by Gene Program Negotiation (GPN) are conserved beyond gastric cancer, we applied the complete analytical pipeline without modification to an independent glioblastoma (GBM) single-cell RNA sequencing dataset (GSE131928). Only GBM-specific biological gene programs were substituted, whereas graph construction, Gene Program Coherence Index (GPCI) calculation, and regulatory state classification were performed using the identical computational framework established in the discovery cohort. Conventional dominant-program assignment identified the major GBM transcriptional programs, including AC-like, MES-like, NPC-like, OPC-like, endothelial, hypoxia, interferon, and proliferation programs (**Fig. 6A**). Application of GPN to the same cells revealed five regulatory decision states—Competition, Consensus, Dominance, Negotiation, and Low activity—that were broadly distributed throughout the transcriptional landscape rather than being restricted to individual dominant programs (**Fig. 6B**). This observation indicates that regulatory organization represents an additional layer of cellular heterogeneity beyond conventional program assignment. Consistent with our observations in gastric cancer, each dominant biological program contained multiple GPN-derived regulatory states (**Fig. 6C**). Cells sharing the same dominant transcriptional program were distributed across Consensus, Competition, Negotiation, Dominance, and Low activity states, demonstrating that dominant program identity alone does not uniquely determine the underlying regulatory organization of individual cells. These findings further support the notion that GPN captures regulatory interactions among simultaneously active biological programs rather than simply identifying the strongest transcriptional signature. We next examined local regulatory conflict across GPN-derived decision states. As observed in the discovery dataset, Competition cells exhibited substantially higher local program conflict than all other regulatory states, whereas Consensus cells consistently maintained low levels of antagonistic regulatory interactions (**Fig. 6D**). This reproducible pattern indicates that elevated local conflict represents a conserved feature of Competition states across distinct tumor types.

**Figure 6.**
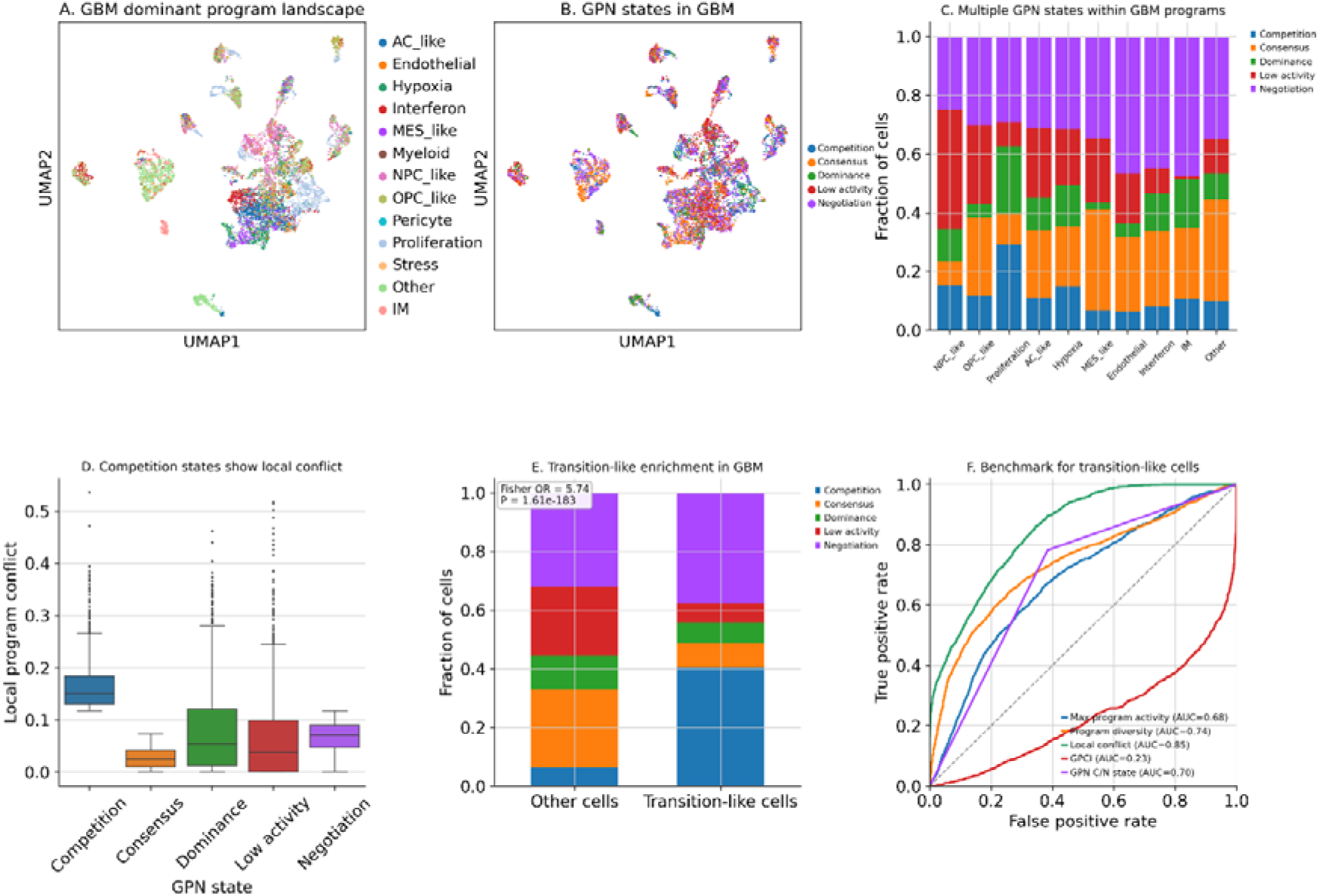
Independent validation of the Gene Program Negotiation framework in glioblastoma single-cell transcriptomes. To evaluate the generalizability of the proposed Gene Program Negotiation (GPN) framework, the complete analytical pipeline was independently applied to the publicly available glioblastoma (GBM) single-cell RNA sequencing dataset (GSE131928) without modifying the core algorithmic framework. Only GBM-specific biological gene programs were substituted, whereas graph construction, Graph-based Gene Program Coherence Index (GPCI), and regulatory decision-state classification remained identical to those used in gastric cancer. (A) UMAP visualization of GBM single-cell transcriptomes colored according to the dominant biological gene program assigned to each cell using conventional independent gene-program scoring. Cells are annotated according to the highest-scoring biological program, including AC-like, MES-like, NPC-like, OPC-like, endothelial, immune, hypoxia, proliferation, stress, and additional GBM-associated programs. (B) UMAP visualization of the same cells colored according to GPN-derived regulatory decision states. Cells were classified into Competition, Consensus, Dominance, Negotiation, or Low activity using graph-derived regulatory coherence, local regulatory conflict, regulatory dominance, and overall biological program activity. (C) Distribution of GPN-derived regulatory decision states within each dominant biological program. Multiple regulatory decision states are observed within every dominant program category, demonstrating that cells sharing the same dominant transcriptional program nevertheless exhibit distinct regulatory organizations captured by GPN. (D) Local regulatory conflict across GPN-derived regulatory decision states. Competition cells exhibit substantially elevated local regulatory conflict compared with the remaining regulatory states, whereas Consensus cells display consistently low antagonistic regulatory interactions, supporting the biological interpretation of the inferred regulatory decision states. (E) Enrichment of GPN-derived regulatory decision states among transition-like GBM cells. Transition-like cells, operationally defined by elevated local regulatory conflict, sustained multi-program activity, and intermediate regulatory coherence, are significantly enriched for Competition and Negotiation states relative to the remaining cell population (Fisher’s exact test, odds ratio = 5.74, P = 1.61 × 10□¹□³), indicating that GPN identifies regulatory organizations associated with dynamic cellular plasticity in an independent cancer type. (F) Receiver operating characteristic (ROC) analysis comparing GPCI with individual regulatory metrics, including maximum program activity, program diversity, local regulatory conflict, and GPN state classification, for identifying transition-like cells. GPCI achieved the highest predictive performance, demonstrating that integrating graph coherence, regulatory activity, and program diversity provides superior discrimination of transition-like cells compared with individual regulatory metrics.

Finally, we investigated whether regulatory decision states were associated with transcriptional plasticity in GBM. Transition-like cells were significantly enriched for Competition and Negotiation states compared with the remaining cell population (Fisher’s exact test, odds ratio = 5.74, *P* = 1.61 × 10□¹□³; **Fig. 6E**). This enrichment closely mirrors the results obtained in gastric cancer and demonstrates that regulatory competition preferentially occurs in cells undergoing dynamic state transitions. Collectively, these findings indicate that the GPN framework identifies conserved patterns of regulatory decision-making across biologically distinct malignancies, supporting the robustness and general applicability of the proposed computational framework.

To assess the robustness of the proposed GPN framework, classification thresholds were systematically varied over a broad range of parameter values. Despite these perturbations, the overall proportions of GPN-derived regulatory decision states, cell-level state assignments, and the enrichment of Competition and Negotiation states among transition-like cells remained highly consistent (**Supplementary Figs. S1–S3**). These findings indicate that the proposed framework is robust to moderate variations in threshold selection and does not rely on a single empirical parameter configuration.

## Discussion

Single-cell transcriptomics has greatly expanded our ability to characterize cellular heterogeneity by identifying transcriptionally distinct cell populations and quantifying the activity of biological gene programs. Nevertheless, most current analytical frameworks implicitly assume that cellular identity can be adequately represented by independent measurements of individual biological programs. This assumption overlooks an important aspect of cellular regulation, namely that multiple gene programs are frequently active simultaneously and collectively influence cellular behavior through cooperative and antagonistic interactions.

Here, we introduce Gene Program Negotiation (GPN), a graph-based computational framework that explicitly models regulatory decision-making among concurrently active biological programs. Rather than focusing solely on the magnitude of program activity, GPN reconstructs cell-specific regulatory interaction networks and quantifies how active programs cooperate, compete, or dominate one another within the local transcriptional context. This approach provides a complementary layer of biological information that is not captured by conventional gene-program scoring alone.

A central finding of this study is that dominant biological programs do not uniquely determine cellular regulatory organization. Cells assigned to the same dominant transcriptional program frequently occupied distinct GPN-derived regulatory states, including Consensus, Competition, Negotiation, Dominance, and Low activity. Similarly, regulatory coherence varied substantially among cells sharing comparable levels of program activity. These observations indicate that transcriptional identity is determined not only by which biological programs are active but also by how these programs interact within individual cells. Consequently, GPN introduces a previously underexplored dimension of single-cell organization that complements existing cell-state annotations.

An important conceptual aspect of the proposed framework is the explicit definition of the Negotiation state. Rather than representing an undefined intermediate group, Negotiation corresponds to a biologically interpretable regulatory configuration characterized by concurrent activity of multiple biological programs without complete agreement or dominance. Such cells maintain balanced regulatory influences while exhibiting intermediate levels of regulatory coherence and local conflict, suggesting that regulatory negotiation itself represents a distinct mode of cellular organization. This interpretation extends conventional views of cell identity by emphasizing how multiple active biological programs are integrated rather than simply identifying the single most active program. An additional strength of the proposed framework is its ability to identify regulatory competition associated with cellular plasticity. Across both gastric cancer and an independent glioblastoma dataset, Competition states consistently exhibited elevated local regulatory conflict and were significantly enriched among transition-like cells. These findings suggest that regulatory competition may represent a conserved feature of dynamic cell-state remodeling rather than a tumor-specific phenomenon. Importantly, these associations emerged without incorporating trajectory information into the GPN algorithm itself, indicating that local regulatory organization independently captures aspects of cellular plasticity. Unlike trajectory inference, RNA velocity, or lineage reconstruction methods, GPN is not designed to estimate temporal progression or developmental directionality. Instead, it quantifies the regulatory organization underlying the instantaneous transcriptional state of each individual cell. The conceptual relationship between GPN and representative single-cell analytical frameworks is summarized in Table 1.

**Table 1.**
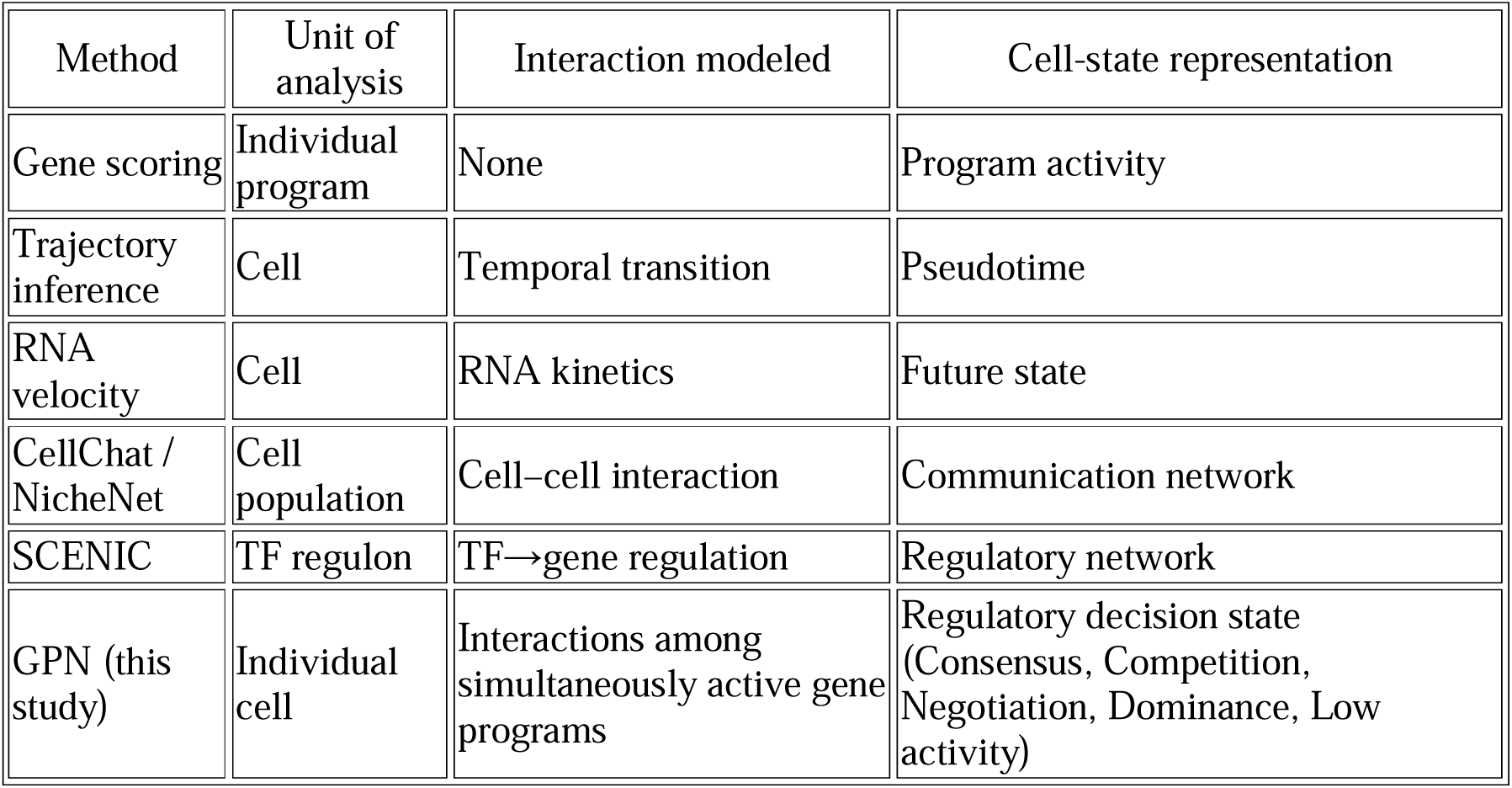
Comparison of Gene Program Negotiation with existing single-cell analytical frameworks.

Consequently, GPN should be viewed as complementary to existing single-cell analytical approaches rather than as a replacement. Integrating GPN with trajectory inference, spatial transcriptomics, perturbation experiments, or lineage tracing may provide a more comprehensive understanding of how regulatory interactions evolve during cellular transitions.

From a computational perspective, the proposed framework is intentionally modular. Because GPN operates on gene-program activity matrices rather than raw expression values, the method can be combined with diverse gene-program estimation approaches, including curated signatures, non-negative matrix factorization, topic models, or latent-factor representations. Similarly, although local transcriptional neighborhoods were defined using conventional k-nearest-neighbor graphs in this study, alternative neighborhood definitions or graph-learning strategies could readily be incorporated without altering the overall framework. This flexibility may facilitate application across diverse biological systems and future computational developments.

Several limitations should also be acknowledged. First, regulatory interactions were inferred from local transcriptional similarity and therefore represent statistical associations rather than direct molecular regulatory mechanisms. Experimental perturbation studies will be required to determine whether the inferred cooperative and antagonistic relationships correspond to causal regulatory interactions. Second, classification of regulatory decision states relies on empirically selected thresholds for regulatory activity, conflict, and dominance. Although identical parameters generalized across independent cancer datasets in the present study, adaptive thresholding or probabilistic state assignment may further improve robustness across broader biological contexts. Finally, the current study focused primarily on transcriptomic regulatory programs. Extending GPN to multimodal single-cell datasets incorporating chromatin accessibility, proteomics, or spatial information may enable more comprehensive characterization of regulatory decision-making.

Overall, this study introduces a conceptual framework in which cellular identity is viewed not simply as the activation of individual biological programs but as the outcome of regulatory negotiation among multiple simultaneously active processes. By explicitly modeling regulatory coherence, competition, and dominance, GPN provides a quantitative description of regulatory decision-making that extends beyond conventional gene-program analysis. We anticipate that this framework will be broadly applicable across diverse biological systems and may contribute to understanding cellular plasticity, disease progression, and therapeutic response at single-cell resolution.

## METHODS

### Study design

Tumor cells frequently activate multiple transcriptional programs simultaneously, including stemness, epithelial–mesenchymal transition (EMT), hypoxia, stress responses and proliferation.^33,34^ Existing single-cell analytical frameworks generally summarize these complex regulatory landscapes by assigning each cell to a single dominant transcriptional program.^34^ Although effective for cell-type annotation, such approaches do not explicitly model interactions among concurrently active biological programs and therefore cannot distinguish coordinated regulatory activation from transcriptional competition.

To address this limitation, a graph-based computational framework termed Gene Program Negotiation (GPN) was developed to quantify local interactions among biological programs at single-cell resolution. Instead of representing each cell by its strongest transcriptional program, GPN reconstructs a cell-specific program interaction network using the local transcriptional neighborhood of each cell.^31,35–37^ The resulting graph enables simultaneous quantification of regulatory coherence, antagonism and program diversity, allowing regulatory decision states to be identified directly from transcriptional organization.

The analytical workflow consisted of four sequential steps: (i) estimation of biological program activities, (ii) reconstruction of cell-specific program interaction graphs, (iii) calculation of graph-derived regulatory metrics, including the Graph-based Gene Program Coherence Index (GPCI), and (iv) classification of cells into regulatory decision states followed by biological interpretation and independent cross-cancer validation.

### Single-cell RNA sequencing datasets

The discovery analysis was performed using the publicly available human gastric cancer single-cell RNA sequencing dataset GSE184198,^38^ obtained from the Gene Expression Omnibus (GEO). This dataset contains malignant epithelial cells together with stromal and immune populations isolated from primary gastric cancer specimens, thereby providing a comprehensive representation of the tumor microenvironment. To evaluate the generalizability of the proposed framework, an independent glioblastoma single-cell RNA sequencing dataset (GSE131928) was analyzed using an identical computational pipeline.^39^ No parameters were modified between discovery and validation analyses, allowing assessment of the robustness of the framework across biologically distinct tumor types.

### Preprocessing of single-cell transcriptomes

Raw count matrices were imported into Scanpy (v1.10).^40^ Cells expressing fewer than 200 genes were excluded. Genes detected in fewer than three cells were removed. Cells exhibiting excessive mitochondrial transcript abundance (>25%) were discarded to minimize technical artifacts associated with low-quality libraries.

Expression counts were normalized to 10,000 transcripts per cell and transformed using the natural logarithm following addition of a pseudocount of one. Highly variable genes were identified using the Seurat variance model, and the 3,000 most variable genes were retained for downstream analyses. ^41^ Gene-expression values were standardized to unit variance before principal component analysis (PCA). Cell–cell neighborhood graphs were constructed using 15 nearest neighbors within the first 30 principal components, followed by Leiden community detection and Uniform Manifold Approximation and Projection (UMAP) visualization.^42^

### Definition of biological gene programs

Biological programs were defined using curated marker genes representing conserved regulatory processes associated with gastric cancer progression.^43–45^ (**Supplementary Table S1**). Programs included epithelial lineage identity, stemness, epithelial–mesenchymal transition (EMT), hypoxia, stress response, proliferation, interferon signaling, immune activation, regulatory T-cell activity, macrophage polarization and fibroblast activation.^45^ Program activity was quantified for every cell using Scanpy’s score_genes algorithm, which estimates the relative enrichment of predefined gene signatures while correcting for gene-expression background.^41^ Only genes detected within each dataset were included during score estimation.

### Cell-specific regulatory interaction graphs

For every individual cell, a local transcriptional neighborhood was defined using the k-nearest-neighbor graph in PCA space.^40^ Program activity vectors were standardized across all cells before local program–program correlations were estimated within each neighborhood. These local correlation matrices describe regulatory compatibility among biological programs under the local transcriptional context.^46^

To incorporate the magnitude of program activation, positive program activities were transformed into pairwise activation weights using an outer-product formulation. The final cell-specific interaction graph was generated through element-wise multiplication of local correlation matrices and activation-weight matrices. Consequently, every cell was represented by a weighted graph describing cooperative and antagonistic interactions among simultaneously active biological programs.

### Graph-based Gene Program Coherence Index (GPCI)

To summarize the overall organization of each cell-specific regulatory negotiation network, we introduced the Graph-based Gene Program Coherence Index (GPCI), a graph-derived metric that provides a quantitative summary of network-wide regulatory organization. For each cell,

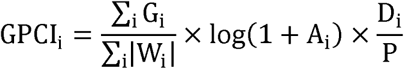

where G_i_ denotes the weighted regulatory interaction graph,^47^ W_i_ represents activation weights, A_i_ corresponds to the total biological program activity, D_i_ denotes effective program diversity, and *P* represents the total number of biological programs.

The GPCI was formulated to quantify coordinated regulatory organization arising from the simultaneous activation of multiple biological programs. Its first component measures graph coherence, representing the overall agreement among regulatory interactions within the cell-specific negotiation network.^48^ The second component reflects the total regulatory signal strength and is logarithmically transformed to reduce the influence of extremely active programs while preserving relative differences in regulatory activity. The final component measures effective program diversity using Shannon entropy, thereby emphasizing coordinated participation of multiple biological programs rather than activation of a single dominant process. Collectively, these three components were combined multiplicatively so that high GPCI values are achieved only when regulatory activity is simultaneously strong, broadly distributed, and mutually coordinated. Consequently, reduction in any one component proportionally decreases the overall GPCI, reflecting disruption of coordinated regulatory organization.

The multiplicative formulation was intentionally adopted because coordinated regulatory organization requires simultaneous agreement, sufficient regulatory activity, and participation of multiple biological programs. An additive formulation would permit high scores despite deficiency in one component, whereas the multiplicative formulation requires all three regulatory properties to be simultaneously satisfied.

Higher GPCI values indicate globally coordinated transcriptional organization, whereas lower values reflect increasing regulatory disagreement among simultaneously active programs.

### Local regulatory conflict

Regulatory competition was quantified using the fraction of negative interactions within each local program interaction graph,^49^

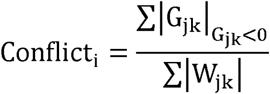

where larger values indicate stronger antagonistic relationships among active biological programs.^50,51^ Unlike GPCI, which summarizes global regulatory coordination, the conflict score specifically measures local regulatory incompatibility.^52^

### Program diversity

To quantify regulatory complexity, effective program diversity was estimated using the exponential form of Shannon entropy, 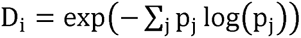 where p_j_ represents the normalized contribution of biological program j. This formulation estimates the effective number of simultaneously active biological programs within each cell while reducing sensitivity to weakly expressed programs. ^53^ ^54^

### Identification of Gene Program Negotiation states

Regulatory decision states were identified by integrating graph coherence, local conflict, overall program activity, regulatory dominance, and effective program diversity.^55^ ^56^ Cells with total program activity in the lowest 20% of the distribution were classified as Low activity. ^4^ Cells in which the dominant program accounted for more than 62% of total program activity were classified as Dominance. Cells with local regulatory conflict in the highest 20% of the distribution were assigned to Competition, whereas Consensus required GPCI above the 70th percentile together with local conflict below the 60th percentile. Cells that did not satisfy the criteria for Low activity, Dominance, Competition, or Consensus were assigned to the Negotiation state. Conceptually, Negotiation states correspond to cells exhibiting concurrent activation of multiple biological programs without clear dominance or complete regulatory agreement. These cells occupy an intermediate regulatory regime characterized by balanced graph coherence and local regulatory conflict while maintaining substantial multi-program activity.^28^ Accordingly, the Negotiation state represents a biologically interpretable regulatory configuration arising from balanced interactions among multiple active biological programs rather than simply representing cells that do not satisfy the other state definitions. Importantly, these classifications emerged directly from graph-derived regulatory properties rather than predefined biological annotations. ^36^

### Trajectory analysis

Diffusion pseudotime was estimated using Scanpy to investigate dynamic changes in regulatory organization during tumor-cell progression. Transition-like regulatory scores were calculated by integrating standardized conflict, total program activity, effective program diversity and GPCI values, thereby identifying cells undergoing active regulatory reorganization.

### Biological interpretation

Average biological program activities were calculated for each regulatory decision state. Enrichment of highly active programs was quantified using log□odds ratios, whereas associations between transition-like cells and regulatory decision states were evaluated using Fisher’s exact test. Together, these analyses connected graph-derived regulatory organization with interpretable biological phenotypes.

### Cross-cancer validation

The complete analytical pipeline was independently applied to the glioblastoma dataset (GSE131928) without modification of algorithmic parameters. Only lineage-specific gene programs were replaced with glioblastoma-relevant regulatory programs while preserving all graph construction procedures, (**Supplementary Table S2**) graph-derived metrics and state-classification criteria. This strategy enabled assessment of whether GPN captures conserved principles of transcriptional organization across biologically distinct malignancies.

### Statistical analysis

Statistical analyses were performed using Python together with the Scanpy, NumPy, SciPy and Pandas libraries. Differences among regulatory decision states were evaluated using the Kruskal–Wallis test. Correlation analyses were performed using Spearman’s rank correlation coefficient, and categorical enrichment was assessed using Fisher’s exact test. All statistical tests were two-sided, and statistical significance was defined as *P*<0.05.

## Declarations

## Ethics approval and consent to participate

N/A

## Consent for publication

N/A

## Availability of data and materials

The analysis codes and datasets used and/or generated during the current study are available from Dr. Ji-Yong Sung upon reasonable request. Interested researchers may contact Dr. Sung via email at to obtain access to the relevant materials.

## Competing interests

The authors declare no interests.

## AI Use Declaration

AI tools were used only for English grammar correction and language polishing.

## Funding

This research was supported by a grant of Korean ARPA-H Project through the Korea Health Industry Development Institute (KHIDI), funded by the Ministry of Health & Welfare, Republic of Korea (grant number: RS-2025-25456722) and supported by the “Regional Innovation Systems & Education (RISE)” through the Seoul RISE Center, funded by the Ministry of Education (MOE) and the Seoul Metropolitan Government. (2026-RISE-01-022-05).

## Authors’ contributions

Conceptualization & Investigation: JYS, JHC; Methodology: JYS; Data analysis: JYS; Writing-original draft: JYS; Writing-review & editing: JYS, JHC; Supervision: JYS, JHC; Project administration: JYS, JHC; Funding acquisition: JHC; Interpretation of the results; JYS, JHC. All authors have read and agreed to the published version of the manuscript.

## Acknowledgements

N/A

## Notes

### Competing Interest Statement

The authors have declared no competing interest.

