## Supplementary Figure 1 for "Gene Program Negotiation Defines Cellular Identity in Single-Cell Transcriptomes"

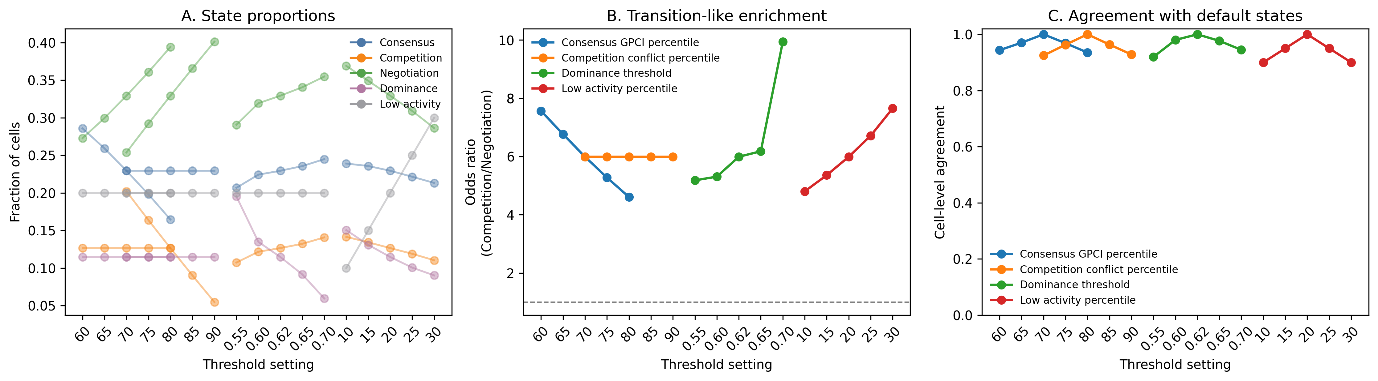

**Supplementary Fig. S1. Robustness of GPN-derived regulatory decision state classification across alternative threshold settings.**

(A) Relative proportions of Consensus, Competition, Negotiation, Dominance, and Low activity states obtained after systematic variation of the classification thresholds for Consensus (GPCI percentile), Competition (local conflict percentile), Dominance (dominance score), and Low activity (program activity percentile). The overall distribution of regulatory decision states remained largely stable across all threshold settings.

(B) Odds ratios describing enrichment of Competition and Negotiation states among transition-like cells under alternative threshold settings. Similar enrichment patterns were observed across all tested parameter values, indicating that the principal biological conclusions were not sensitive to the specific threshold selection.

(C) Cell-level agreement between the default GPN classification and classifications obtained using alternative threshold settings. Agreement exceeded 90% under all tested conditions, supporting the robustness and reproducibility of the proposed GPN framework.
