## Supplementary Figure 2 for "Gene Program Negotiation Defines Cellular Identity in Single-Cell Transcriptomes"

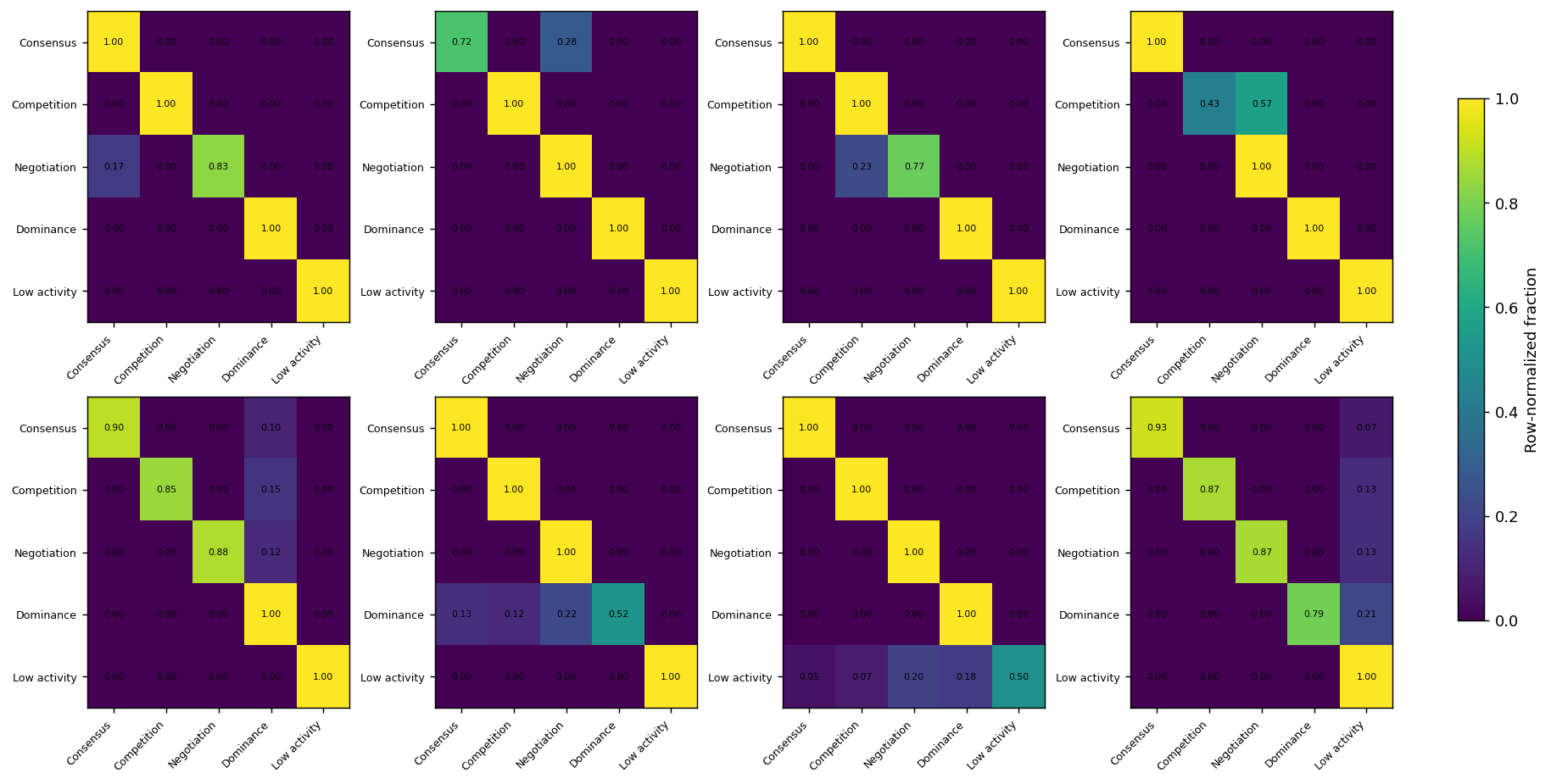

**Supplementary Fig. S2. Cell-level agreement between default and alternative GPN classifications.**

Heatmaps showing row-normalized confusion matrices comparing the default GPN-derived regulatory decision states with classifications obtained after varying the thresholds for Consensus (GPCI percentile), Competition (local conflict percentile), Dominance (dominance score), and Low activity (program activity percentile). Rows correspond to regulatory decision states assigned using the default parameters, and columns correspond to classifications obtained using alternative threshold values. High values along the diagonal indicate that the majority of cells retained their original regulatory decision state despite threshold variation, whereas off-diagonal elements represent reassigned cells. These results demonstrate that the proposed GPN framework produces highly reproducible regulatory state assignments across a broad range of threshold settings.
