## Supplementary Figure 3 for "Gene Program Negotiation Defines Cellular Identity in Single-Cell Transcriptomes"

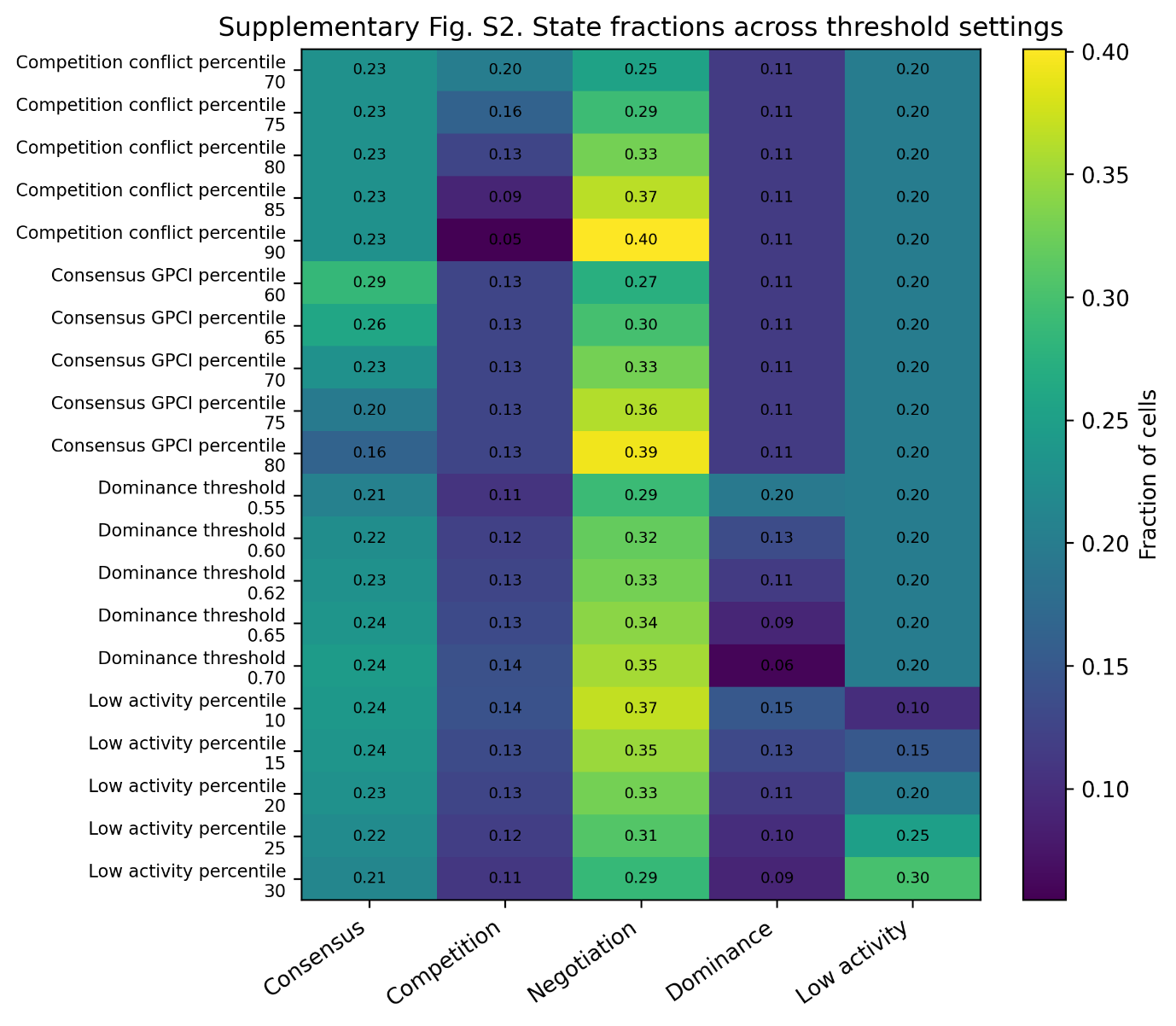

**Supplementary Fig. S3. Stability of GPN-derived regulatory decision state proportions across alternative threshold settings.**

Heatmap summarizing the fraction of cells assigned to each GPN-derived regulatory decision state following systematic variation of the classification thresholds for Consensus (GPCI percentile), Competition (local conflict percentile), Dominance (dominance score), and Low activity (program activity percentile). Rows represent individual threshold settings, whereas columns represent the five regulatory decision states. Cell colors and numerical values indicate the fraction of cells assigned to each state. Across all tested parameter settings, the overall distribution of Consensus, Competition, Negotiation, Dominance, and Low activity states remained highly consistent, supporting the robustness of the GPN framework to moderate changes in classification thresholds.
