## Supplementary Tables for "Gene Program Negotiation Defines Cellular Identity in Single-Cell Transcriptomes"

**Supplementary Table S1. Glioblastoma-specific gene programs used for cross-cancer validation**

| Gene Program | Biological Function | Marker Genes |
| --- | --- | --- |
| Epithelial | Gastric epithelial identity | EPCAM, KRT8, KRT18, KRT19, MUC1, CLDN3, CLDN4, TACSTD2 |
| Gastric lineage | Gastric epithelial differentiation | MUC5AC, MUC6, TFF1, TFF2, PGC, GKN1, GKN2, GIF |
| Stemness | Cancer stem-cell program | LGR5, PROM1, SOX2, OLFM4, ASCL2, BMI1, CD44, ALDH1A1 |
| EMT | Epithelial–mesenchymal transition | VIM, FN1, ZEB1, ZEB2, SNAI1, SNAI2, TWIST1, ITGA5 |
| Hypoxia | Cellular hypoxia response | HIF1A, VEGFA, CA9, LDHA, SLC2A1, ENO1, PGK1, BNIP3 |
| Stress response | Immediate early and stress-response genes | FOS, JUN, DUSP1, HSPA1A, HSP90AA1, ATF3, IER2, EGR1 |
| Proliferation | Cell-cycle progression | MKI67, TOP2A, PCNA, TYMS, STMN1, UBE2C, BIRC5, CENPF |
| Interferon signaling | Type I interferon response | ISG15, IFIT1, IFIT2, IFIT3, MX1, OAS1, STAT1, IRF7 |
| Immune activation | Adaptive immune activation | PTPRC, CD3D, CD3E, NKG7, GZMB, CD74, HLA-DRA, CXCL13 |
| Regulatory T cell (Treg) | Immunosuppressive T-cell program | FOXP3, IL2RA, CTLA4, TIGIT, IKZF2, TNFRSF18, CCR8 |
| Macrophage | Myeloid/macrophage activation | LYZ, C1QA, C1QB, C1QC, CD68, AIF1, FCGR3A, LST1 |
| Fibroblast | Stromal activation | COL1A1, COL1A2, DCN, LUM, COL3A1, ACTA2, TAGLN, PDGFRB |

**Supplementary Table S2. Glioblastoma-specific gene programs used for cross-cancer validation**

| Gene Program | Biological Function | Marker Genes |
| --- | --- | --- |
| NPC-like | Neural progenitor-like | SOX4, SOX11, DLL3, ASCL1, HES6, TCF12, TUBB3, DCX |
| OPC-like | Oligodendrocyte progenitor-like | PDGFRA, OLIG1, OLIG2, SOX10, CSPG4, BCAN, NKX2-2, PLP1 |
| AC-like | Astrocyte-like | GFAP, AQP4, SLC1A3, ALDOC, CLU, APOE, FABP7, GJA1 |
| MES-like | Mesenchymal-like | CHI3L1, CD44, VIM, SERPINE1, TGFBI, ANXA2, LGALS1, FN1 |
| Proliferation | Cell-cycle progression | MKI67, TOP2A, PCNA, TYMS, STMN1, UBE2C, BIRC5, CENPF |
| Hypoxia | Hypoxia response | HIF1A, VEGFA, CA9, LDHA, SLC2A1, ENO1, PGK1, BNIP3 |
| Stress | Cellular stress response | FOS, JUN, DUSP1, HSPA1A, HSP90AA1, ATF3, IER2, EGR1 |
| Interferon | Interferon signaling | ISG15, IFIT1, IFIT2, IFIT3, MX1, OAS1, STAT1, IRF7 |
| Immune activation | Adaptive immune activation | PTPRC, CD3D, CD3E, NKG7, GZMB, CD74, HLA-DRA, CXCL13 |
| Myeloid | Myeloid lineage | LYZ, C1QA, C1QB, C1QC, CD68, AIF1, FCGR3A, LST1 |
| Endothelial | Vascular endothelial cells | PECAM1, VWF, KDR, FLT1, RAMP2, ESAM, CLDN5, ENG |
| Pericyte | Perivascular stromal cells | RGS5, PDGFRB, ACTA2, TAGLN, MCAM, CSPG4, COL1A1, COL3A1 |
